# Local density shapes complete brood failure across species boundaries in two sympatric songbirds

**DOI:** 10.64898/2026.07.02.734466

**Authors:** Gregory F Albery, Sarah CL Knowles, Carys V Jones, Ben C Sheldon, Josh A Firth

**Affiliations:** School of Natural Sciences, Trinity College Dublin, Dublin, Republic of Ireland; Department of Biology, Georgetown University, Washington, DC; Department of Biology, University of Oxford, UK; Edward Grey Institute, Dept of Biology, University of Oxford, UK; School of Biology, University of Leeds, UK

## Abstract

Reproduction in species with parental care involves sustaining a brood of offspring through an energetically demanding period, when shifts in resource availability, weather, predation risk, and parental condition can strongly alter offspring survival. The most extreme outcome is complete brood failure (death of all offspring), which is relatively frequent in many bird species and may occur when conditions cross a viability threshold. Although complete brood failure is important for shaping fitness variation and population dynamics, we have limited understanding of how intra– and interspecific density dependence governs these events, or of how factors such as habitat quality and disease burden contribute to them, because deriving this requires fine-scale, individual-level data collected across generations for multiple overlapping species. Using a dataset totalling 38,509 nesting attempts from great tits (*Parus major*) and blue tits (*Cyanistes caeruleus*) in Wytham Woods, Oxford, UK, we examined how brood failure is shaped by local conspecific and heterospecific density, habitat structure, and avian malaria infection for a subset. Complete brood failure was frequent (14.75%), mostly involving chick mortality in the nest consistent with starvation, rather than brood removal by predators. Relationships between density and brood failure were strong but species-specific. Specifically, great tit failure risk was higher in neighbourhoods that remained densely populated across years, whereas blue tit failure risk was lower where annual great tit or combined density was high, but not where annual blue tit density itself was high. This suggests that local overall density reflects continuing constraint for great tits, while local annual density may partly track favourable within-year conditions and settlement patterns for blue tits. In great tits, failure was also more common where oak density was low and farther from the closest river (Thames), while habitat associations were weak in blue tits. Malaria infection was spatially heterogeneous and covaried with density and habitat, but infection status did not significantly explain complete brood failure. Together, these results show that complete brood failure is shaped by spatially structured local ecological context, and how density dependence in these events can differ in direction and timescale between sympatric species.

## Introduction

For many animal species, reproduction involves not just producing a single offspring at once, but raising a brood of multiple offspring at the same time. This strategy demands substantial parental investment across various aspects of life, including in territory defense, nest construction, and prolonged care of the young (Perrins 1965; Slagsvold 1981). In such species, nestmates share the same local environment and experience (in terms of external conditions and parental care), and often compete with each other for resources, so the loss of each offspring is often non-independent within each brood (Cody 1966). Indeed, all offspring in a brood may die (‘brood failure’), often all around the same time (Perrins 1965) and leaving the parents with no reproductive output from that attempt despite their – often substantial – efforts (Williams 1966). Such whole-brood failures represent extreme fitness costs (Charnov and Krebs 1974) because they entail the complete loss of a breeding attempt (Perrins 1965; Slagsvold 1981) and the loss of all resources invested in that brood (Williams 1966). As such, brood failure can be considered as a particularly consequential event, which by eliminating an entire breeding attempt might contribute disproportionately to variation in reproductive success and have implications for population change (Perrins 1965; Williams 1966; Martin 1995; Sutherland 1996). At the individual level, understanding complete failure may be further biologically informative because it may reflect a rapid transition from an apparently viable brood to collapse, rather than simply the far end of a gradual brood-reduction process (Clark and Wilson 1981; Schöll and Hille 2020).

Complete brood mortality is documented across diverse taxa, sometimes affecting a sizable fraction of nesting attempts, particularly in avian species (Cody 1966; Martin 1995). Understanding the causes of brood failure is therefore important for explaining variation in individual fitness (Gamelon et al. 2021), and may also be useful for gaining insight into how natural selection shapes reproductive strategies to mitigate such risks (Stearns 1992). Indeed, the occurrence of, and variation in, brood failure is likely to be shaped by a combination of diverse individual-level factors (relating to individual quality) and multiple ecological pressures (dependent on the local environment), all of which may interact with demography (such as population density). However, it remains unclear to what extent we can predict the likelihood of brood failure at the individual level, how ecological factors across different local environments may shape this, and whether these effects operate across different species.

The probability of brood failure, as with many other components of an animal’s ability to survive and reproduce, can depend on resource availability (Schöll and Hille 2020; Kirkpatrick, Conway, and Ali 2009), competition (Mennerat et al. 2018), predation (Schöll and Hille 2020; Kirkpatrick, Conway, and Ali 2009; Mudge and Talbot 1993), brood parasitism (Payne 1977), parasite infestation (Hurtrez-Boussès et al. 1997), environmental stress (Kochert, Steenhof, and Brown 2019), and anthropogenic factors such as pollution, hunting, or land use (Milchev and Georgiev 2012; Mudge and Talbot 1993; Matthysen and Adriaensen 1998; Byholm 2005). Often the proximate cause of brood failures is loss of (or abandonment by) a parent, which may prevent the brood from receiving sufficient resources (Santema and Kempenaers 2018; Mudge and Talbot 1993; Thibault, Sanders, and Jodice 2010). Population density can alter local social structure and encounter rates (Beck et al. 2023; Beck et al. 2024). Higher density can also increase competition for resources, parasite exposure, or predation risk, although the direction and strength of these effects vary across systems (Sih 1984; Wang et al. 2009; Hasik et al. 2024; Albery 2022). In particular, when densities are too high and resources insufficient, excess offspring may regularly suffer and die, particularly in income breeder species. However, while the balance of positive and negative density-dependent factors is well understood in the context of population and community ecology, they are rarely examined at the individual level.

Additionally, researchers have not compared and contrasted the effects of local population densities of multiple closely related species to understand how the presence of each species determines patterns of offspring survival, and understanding of brood failure occurrence in multi-species communities (where local density reflects both conspecific crowding and heterospecific interactions) remains limited. Nevertheless, sympatric, closely related species often compete directly for nest sites and food (Minot 1981; Minot and Perrins 1986; Dhondt 2010), and can also influence one another indirectly via shared predators and parasites, and through the social and informational environment in mixed neighbourhoods (Magrath et al. 2015; Firth et al. 2016; Firth et al. 2018; Keen et al. 2020). Yet, it is unclear to what degree intra– and interspecific population density drives brood failure, and whether these effects act similarly across individuals of different species sharing the same environment.

Here we draw on a long-term study of individual-based monitoring of great tits (*Parus major*) and blue tits (*Cyanistes caeruleus*) in Wytham Woods, Oxford, to examine the ecology underpinning complete brood failure in natural populations (Perrins 1965; Gamelon et al. 2021). This represents a particularly strong model for examining brood failure associations with density across species, as great tits are strong interference competitors for nest sites, while blue tits can be effective exploitative competitors for food during breeding, particularly where nest sites are not limiting (Minot 1981; Minot and Perrins 1986; Dhondt 2010). Further, because these species overlap in habitat and phenology and share key food resources, heterospecific density could shape performance even when conspecific density is held constant (Perrins 1965; Dhondt 2010; Gamelon et al. 2021). More broadly, heterospecific density may also track aspects of habitat quality or community composition that are only partly captured by simple habitat metrics, creating the possibility that density-fitness relationships differ in direction or magnitude between interacting species.

As such, we combine breeding bird records spanning almost 60 years from 1965–2023 with fine-scale nest-location, density, habitat, and parasite-prevalence data for both species to investigate how local conspecific and heterospecific densities shape brood failure across time and space, alongside habitat structure and parental infection status. We then discuss how this approach, combining detailed multi-faceted long-term data across two species with spatially explicit Bayesian models (INLA), provides new insights into the causes and consequences of brood failure in wild species (Gokcekus et al.

2023; Woodman et al. 2026).

## Methods

### Study system: the Wytham Woods tits

To understand how population density drives brood failure, we used two cohabiting populations of passerine birds in Wytham Woods, Oxfordshire, UK (51°46’ N, 1°20’ W) (Perrins 1965; Gosler 1993). This study was conducted from 1965 to 2023 using the long-term individual-based study of great tits (*Parus major*) and blue tits (*Cyanistes caeruleus*). For both species, birds breed in nestboxes that are monitored throughout the breeding season. Each breeding attempt consists of obtaining a territory, building a nest within a nestbox, laying and incubating eggs, and rearing offspring. Once hatched, chicks are provisioned by both parents until fledging (usually 17-22 days old). During the breeding season nestboxes are visited to record information on the stages of each breeding attempt (Perrins 1965). Nests were monitored weekly until eggs were found, and thereafter every other day to determine hatch date, or the day that chicks started hatching (day 1). Parents are identified using either capture-and-ringing procedures during the nestling phase and – if previously unringed – are trapped and fitted with a unique BTO (British Trust for Ornithology) metal leg ring or using radio-frequency identification methods (by adding an RFID tag to birds captured from 2007 onwards) allowing passive identification of adults from when nestlings are between 6-14 days old (Firth & Sheldon 2016). At 15 days old, nestlings are ringed and weighed, and mist-netting is carried out during autumn and winter to capture and mark any immigrants and unringed birds.

#### Recording Brood Failure

After ringing and weighing the chicks on day 15, all nests are checked once more after fledging to determine how many chicks left the nest (and the rings of the deceased chicks are recorded). Nestling loss and complete brood failure can happen at any point over the nestling period, and we coded brood failure as 1 if no chicks survived to fledging, and 0 otherwise; we did not include nests with zero hatchlings in the analyses.

Brood failure in this population generally results from abandonment by the parents, either because they cease provisioning the nest or die before rearing is complete (Santema and Kempenaers 2018; Gosler 1993; Jones 2025). To confirm that most complete brood failure events in our dataset occurred because chicks were no longer provisioned (rather than chicks being removed by predators or other causes), we re-examined the original nest-record database for both species (Figs. S1-S3). For each breeding attempt we extracted (i) the binary flags ‘legacy predation’ (used until around 2010), ‘suspected predation’ (used thereafter), and ‘whole brood missing’; (ii) the recorded number of dead chicks left in the nest; and (iii) free-text comments, which we screened for the strings ‘starv*’ or ‘abandon*’. We then compared, for every year, the count and proportion of broods with no fledglings that also carried any predation flag, any starvation/abandonment comment, or at least one dead nestling found in the nest. Across the full time series, predation codes were found in only ∼7% of broods that yielded no fledglings (Jones 2025). Predation codes were most common during the late 1960s–early 1970s but fell to near-zero for most subsequent years (Figs. S1–S2), and whole-brood-missing annotations remained rare throughout (Figs. S1–S3), whereas records of dead chicks in nests were common in the years for which these annotations were consistently available (Fig. S3; but see Julliard et al. 1997). These patterns indicate that the vast majority of complete brood failures are associated with chicks dying in the box, consistent with starvation following parental desertion or death, rather than with predators removing entire broods (Jones 2025). Given the rarity of predation flags and the prevalence of dead chicks in failed nests, we treat ‘no fledglings’ as a reliable proxy for brood failure caused primarily by starvation (Santema and Kempenaers 2018). This assumption is further supported by field observations of emaciated nestlings in failed broods (pers. obs. by all field workers) and by the low incidence of whole-brood disappearance events. We therefore also interpret complete brood failure as a potential welfare-relevant outcome reflecting insufficient parental provisioning, assuming that starving to death carries some stress (Karaer, Čebulj-Kadunc, and Snoj 2023; McCue, Terblanche, and Benoit 2017) which personal observations and previous research (McCue et al. 2017; Karaer et al. 2023) also support.

### Density metrics

We calculated local population density using a previously described pipeline (Gokcekus et al. 2023; Gokcekus et al. 2025), based on the distribution of nesting locations. This approach uses a kernel density estimator that fits a two-dimensional smooth to the distribution of nesting locations. The kernel density estimator produces a two-dimensional spatial distribution of the population’s density, and individuals (nests) are then assigned a local density value based on their location on the kernel. We have used a similar approach to understand ecological and behavioural processes in badgers (Albery et al. 2020) and deer (Hasik et al. 2024), and in meta-analytical contexts (Albery et al. 2020; Hasik et al. 2024; Albery et al. 2025). We carried this out on two timescales: local annual density (fitting a different smooth per year, using each individual’s annual location) and local overall density (fitting a single smooth across the entire study period, using each individual’s lifetime centroid). We also conducted this separately for subsets of birds: great tits only, blue tits only, and great and blue tits combined (“full”). Combining these scales of time and taxonomy therefore gave six measures of density.

### Malaria infection status

We used previously generated PCR-based diagnoses of avian malaria parasite (*Plasmodium* spp.) infection from breeding adult blue tits and great tits in Wytham Woods (Wood et al. 2007; Knowles et al. 2011; Lachish et al. 2011, 2013; Table S1). These malaria data represent a subset of the full breeding dataset: blue tit infection data were available for adults sampled between 2005 and 2010, while great tit infection data were available for adults sampled in 2008 and 2009. Across both host species, blood samples were collected from individually marked breeding adults at a standardised point in the nestling period, typically when broods were 6-14 days post-hatch.

Malaria infections in this system are dominated by two *Plasmodium* morphospecies, *P. relictum* and *P. circumflexum*, which correspond to distinct mitochondrial *cytochrome b* lineages and together comprise the great majority of haemosporidian infections in the Wytham tit populations (Palinauskas et al. 2007; Knowles et al. 2011; Lachish et al. 2011, 2013). We therefore analysed malaria infection status using parasite species-specific binary measures for both host species, giving four host-parasite combinations: great tit-*P. relictum*, great tit-*P. circumflexum*, blue tit-*P. relictum*, and blue tit-*P. circumflexum* (Table S1).

Samples were screened using PCR-based assays described previously for this study system. The primary diagnostic method was a SYBR-green qPCR assay targeting a 188-bp fragment of *Plasmodium cytochrome b*, run in triplicate and scored as positive if any replicate amplified. This qPCR assay also allowed assignment to *P. relictum* or *P. circumflexum* using diagnostic differences in product melting temperature (Knowles et al. 2011). Some blue tit samples were additionally, or previously, screened using nested *cytochrome b* PCR following Waldenström et al. (2004), with parasite species assignment based on sequence identity. Where both diagnostic methods were available, they showed high agreement in overall infection status and in parasite species assignment, although qPCR was generally more sensitive (Knowles et al. 2011; Lachish et al. 2011).

Because our analyses focused on infection prevalence rather than parasitaemia, qPCR output was converted to binary infection status. For each host species and parasite species, individuals were coded as infected or uninfected based on the available diagnostic information. Individuals with mixed *P. relictum* and *P. circumflexum* infections were rare; where present, they were treated as infected in the relevant species-specific analyses for both parasite species. We then linked parental infection status to breeding attempts using parent identities. For brood-failure models, malaria analyses were restricted to breeding attempts for which at least one parent had associated malaria diagnostic data, and infection was coded as present for a given parasite species if either sampled parent was infected. We therefore treated missing malaria diagnoses as missing data, rather than as evidence of absence of infection.

### Analysis

Our final dataset included 38,509 observations spread across a total of 59 years. These comprised 16,956 great tit nesting attempts and 21,553 blue tit nesting attempts. We carried out our analyses with the Integrated Nested Laplace Approximation (INLA), which is a deterministic Bayesian algorithm for fitting linear models (Rue et al. 2009; Lindgren and Rue 2015; Lindgren, Rue, and Lindstrom 2011).

#### Brood failure models

To examine the drivers of brood failure, we fitted two models: one for great tits, and one for blue tits. Brood failure was the binary response variable, with a binomial specification. We fitted year and lay date (day of year) as continuous fixed effects, and nest ID and year as categorical random effects. To explore the spatial determinants of brood failure, we then fitted a series of spatially defined metrics one at a time. These included our six measures of local density (annual and overall; great tit, blue tit, and full). We also included distance from the far northwest corner of the study area, where the Thames river was closest (Thames distance), and local density of oak trees (Oak density; defined in the same way as population density). Because our spatial metrics were highly correlated, we only included one at a time, and kept the best-fitting before attempting to add the remaining variables. We report the minimal models that descriptively fit only one variable at a time, as well as the final model that included only the best-fitting.

#### Malaria models

To examine how local ecological context was associated with malaria prevalence, we fitted malaria infection status as a binary response variable in separate models for each host-parasite combination: great tit-*P. relictum*, great tit-*P. circumflexum*, blue tit-*P. relictum*, and blue tit-*P. circumflexum*. These models included the same spatially distributed explanatory variables used in the brood-failure models, including local annual tit density, distance from the Thames, and oak density. As in the brood-failure analyses, we compared models with and without an SPDE spatial autocorrelation effect to assess whether infection prevalence showed residual spatial structure after accounting for these covariates.

Finally, to test whether malaria infection could explain variation in complete brood failure, we added parental malaria infection status as an explanatory variable to the brood-failure models. These models were fitted separately by host species and parasite species, allowing us to test whether infection with either *P. relictum* or *P. circumflexum* was associated with brood failure in great tits or blue tits. For example, this allowed us to test whether brood failure was more common in high-density areas because higher local density was associated with increased malaria prevalence. Because malaria diagnoses were available only for a subset of breeding attempts, these analyses were restricted to nests for which parental infection data were available.

#### Spatial effects

For all models, after fitting explanatory variables we then fitted a stochastic partial differential equation (SPDE) effect in INLA to examine patterns of spatial heterogeneity and account for spatial autocorrelation (Rue et al. 2009; Lindgren, Rue, and Lindstrom 2011). For each data point, we fitted the X and Y coordinates of the nest, where closer nests in space would be expected to have more similar values for the response variable.

## Results

We discovered strong but contrasting effects of density on brood failure across species. For great tits, local overall density across the study period was associated with greater rates of brood failure, whether considering blue tits (Fig. 1A; Fig. 2A; coeff = 0.17, 95% CrI = 0.11, 0.23, P < 0.001), great tits (Fig. 1A; coeff = 0.15, 95% CrI = 0.09, 0.21, P < 0.001), or both (Fig. 1A; coeff = 0.17, 95% CrI = 0.11, 0.23, P < 0.001). Brood failure in blue tits was negatively associated with local annual density of great tits (Fig. 1B; Fig. 2B; coeff = –0.06, 95% CrI = –0.11, –0.01, P = 0.02) and both species (Fig. 1B; coeff = –0.06, 95% CrI = –0.11, –0.00, P = 0.03), although not with blue tits alone (Fig. 1B; coeff = – 0.04, 95% CrI = –0.10, 0.03, P = 0.26). There were strong positive effects of distance from the Thames (Fig. 1A; coeff = 0.18, 95% CrI = 0.12, 0.24, P < 0.001), and strong negative effects of oak density (Fig. 1A; coeff = –0.16, 95% CrI = –0.22, –0.10, P < 0.001), on great tit brood failure, but these did not manifest for blue tits (Fig. 1B). There was strong evidence for spatial heterogeneity in brood failure in both species: including an SPDE autocorrelation effect substantially improved model fit when modelling great tit brood failure (Fig. 3; ΔDIC = –24.15) and when modelling blue tit brood failure (Fig. 3; ΔDIC = –26.49). Finally, there were strong positive effects of lay date on blue tits (coeff = 0.403, 95% CrI = 0.33, 0.476, P < 0.001) and great tits (coeff = 0.216, 95% CrI = 0.137, 0.296, P < 0.001).

**Figure 1:**
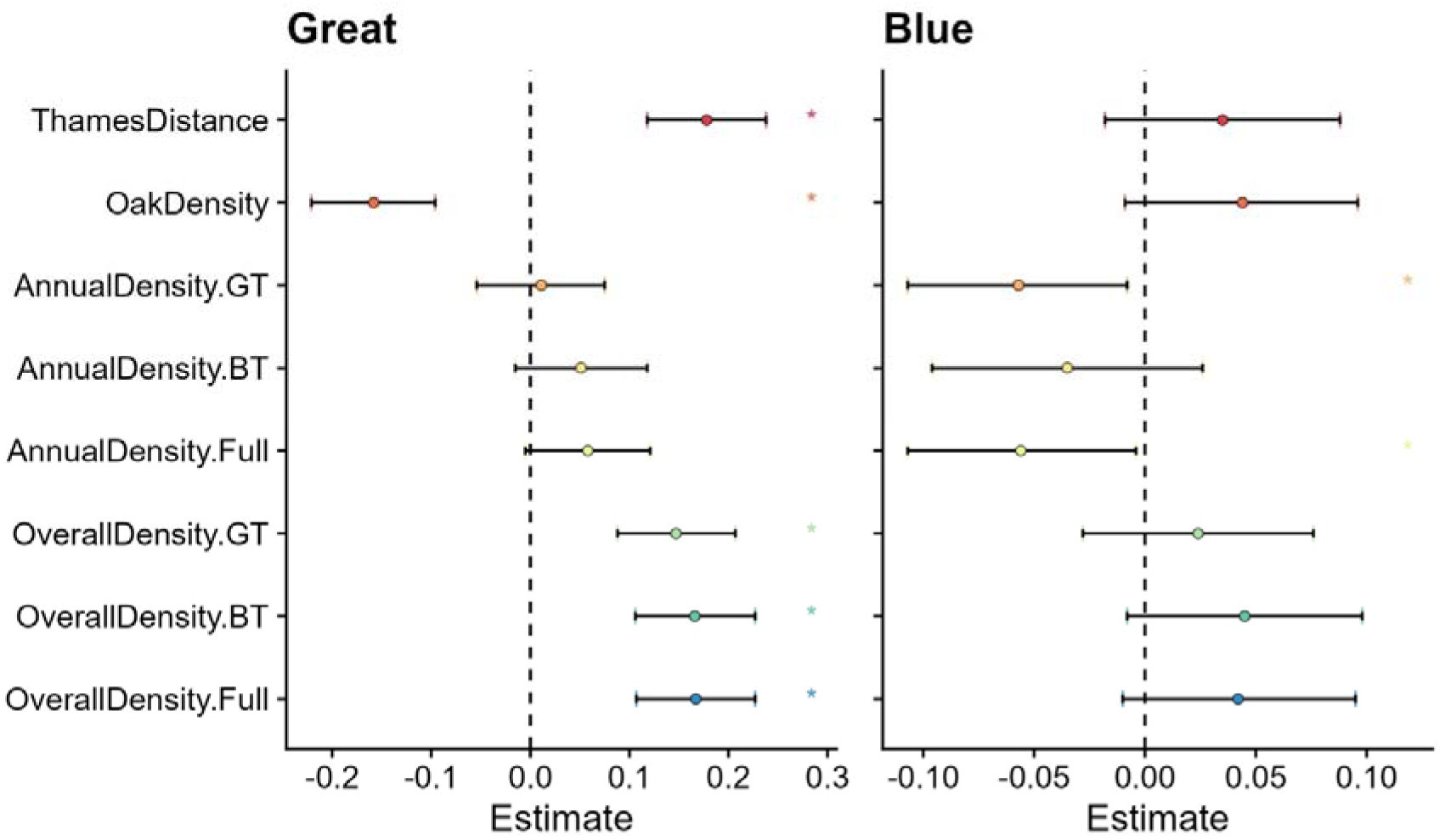
Estimates of spatially varying effects on brood failure in great tits (Panel A) and blue tits (Panel B). Points represent posterior estimates for mean effect sizes, error bars denote 95% credible intervals in standard deviations, and colour denotes individual models with single spatial variables fitted. Asterisks denote that the effect is significant (i.e., credible intervals do not overlap with 0).

**Figure 2:**
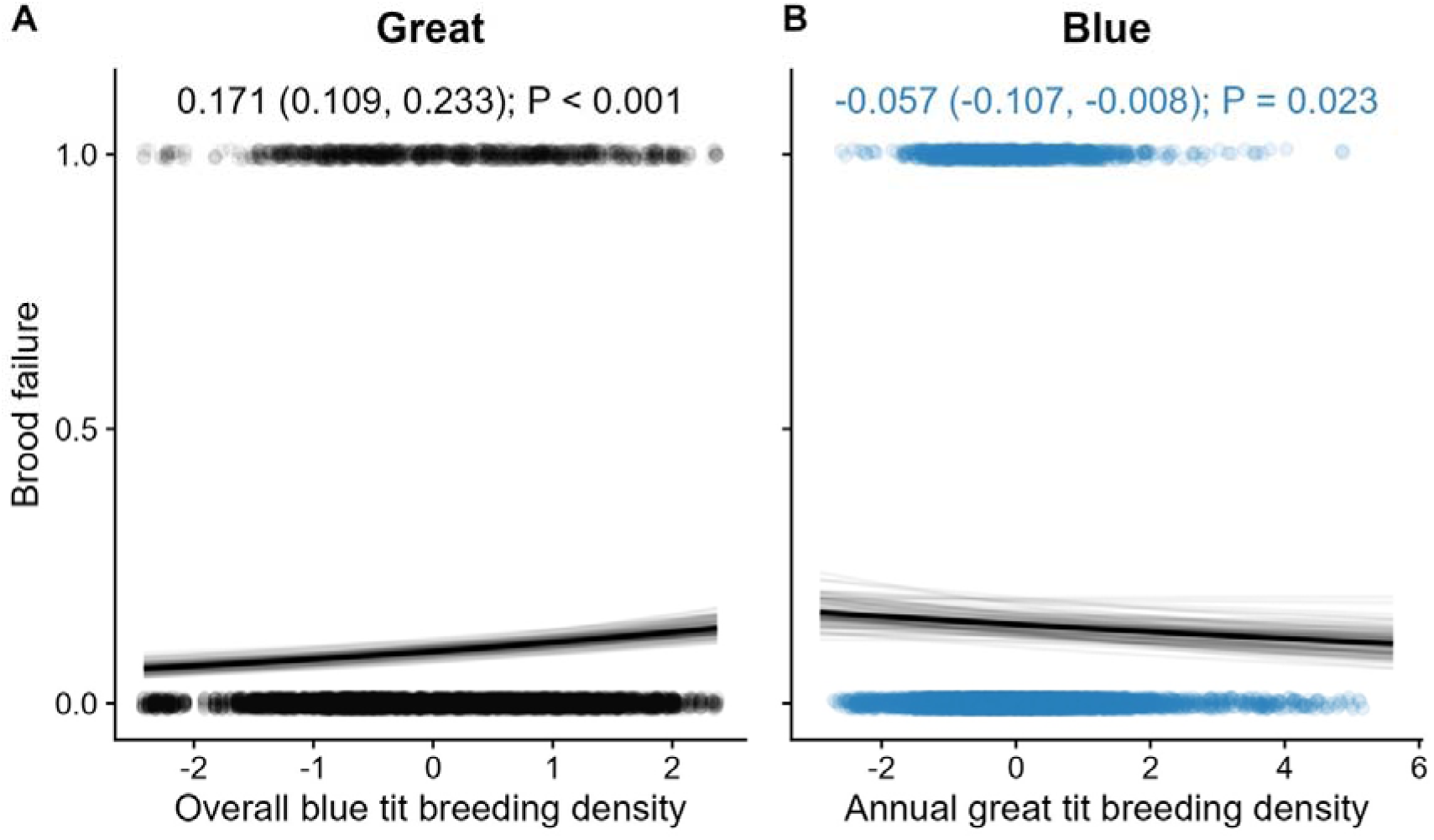
Relationships between local density and brood failure rates in great tits (Panel A) and blue tits (Panel B). The dark black line represents the mean of the posterior distribution for the model estimates; the light grey lines are 100 random draws from the posterior to represent uncertainty. Points denote observations of individual nests, with transparency to allow for visualisation of overplotting.

**Figure 3:**
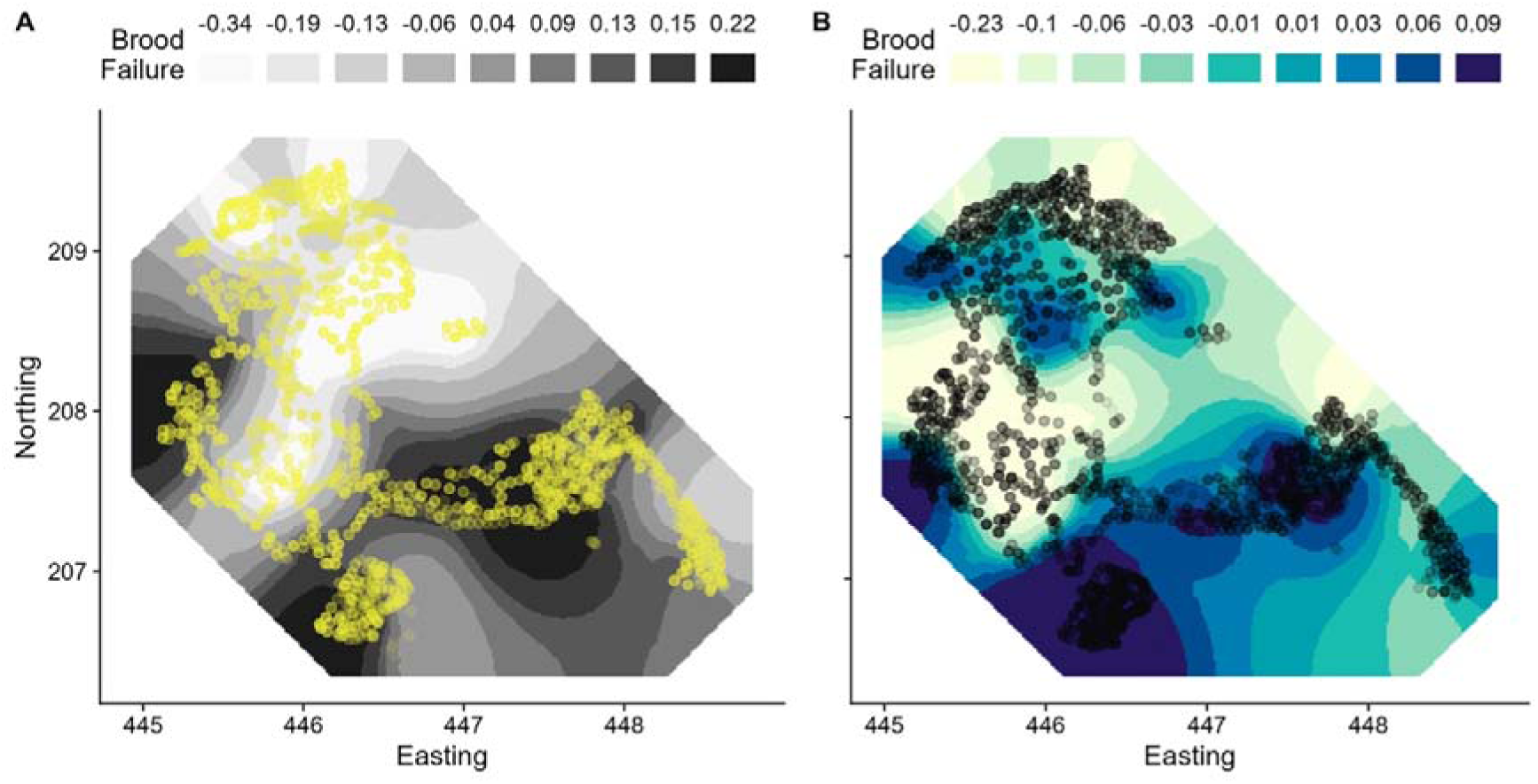
Projected spatial effects describing brood failure rates in great tits (Left) and blue tits (Right). Shown are projections of the spatially-distributed SPDE random effect from the base model, without spatial fixed effects. Shading of the map denotes the lower bounds of nine quantiles of the spatial effects on the link scale, rounded to two decimal places, with darker colors representing higher brood failure probability. Points represent each nesting location. Easting and Northing are in km.

Across the 2 x 2 comparison of host and malaria species, we report the effects from the full models, in which Thames distance, oak density, and local annual great tit density were fitted together (Fig. 4). For P. circumflexum, prevalence decreased with distance from the Thames in both great tits (–0.90 (–1.10, –0.70), P < 0.001) and blue tits (–0.80 (–0.95, –0.66), P < 0.001). Oak density was negatively associated with P. circumflexum prevalence in both great tits (–0.39 (–0.57, –0.20), P < 0.001) and blue tits (–0.34 (–0.47, –0.20), P < 0.001). Local annual great tit density was positively associated with P. circumflexum prevalence in both great tits (0.54 (0.39, 0.69), P < 0.001) and blue tits (0.43 (0.33, 0.52), P < 0.001). For P. relictum, prevalence increased with distance from the Thames in both great tits (0.26 (0.08, 0.44), P = 0.004) and blue tits (0.26 (0.15, 0.36), P < 0.001). Oak density was not associated with P. relictum prevalence in great tits (0.08 (–0.10, 0.25), P = 0.40), but was positively associated with prevalence in blue tits (0.15 (0.04, 0.26), P = 0.01). Local annual great tit density was not associated with P. relictum prevalence in great tits (0.04 (–0.11, 0.18), P = 0.63), but was negatively associated with prevalence in blue tits (–0.18 (–0.27, –0.08), P < 0.001). These infection patterns showed spatial structuring (Fig. 5), with residual spatial autocorrelation clearly improving model fit for great tit P. circumflexum (ΔDIC = –94.14), blue tit P. circumflexum (ΔDIC = –138.71), and blue tit P. relictum (ΔDIC = –21.13), but not great tit P. relictum (ΔDIC = –1.77).

**Figure 4:**
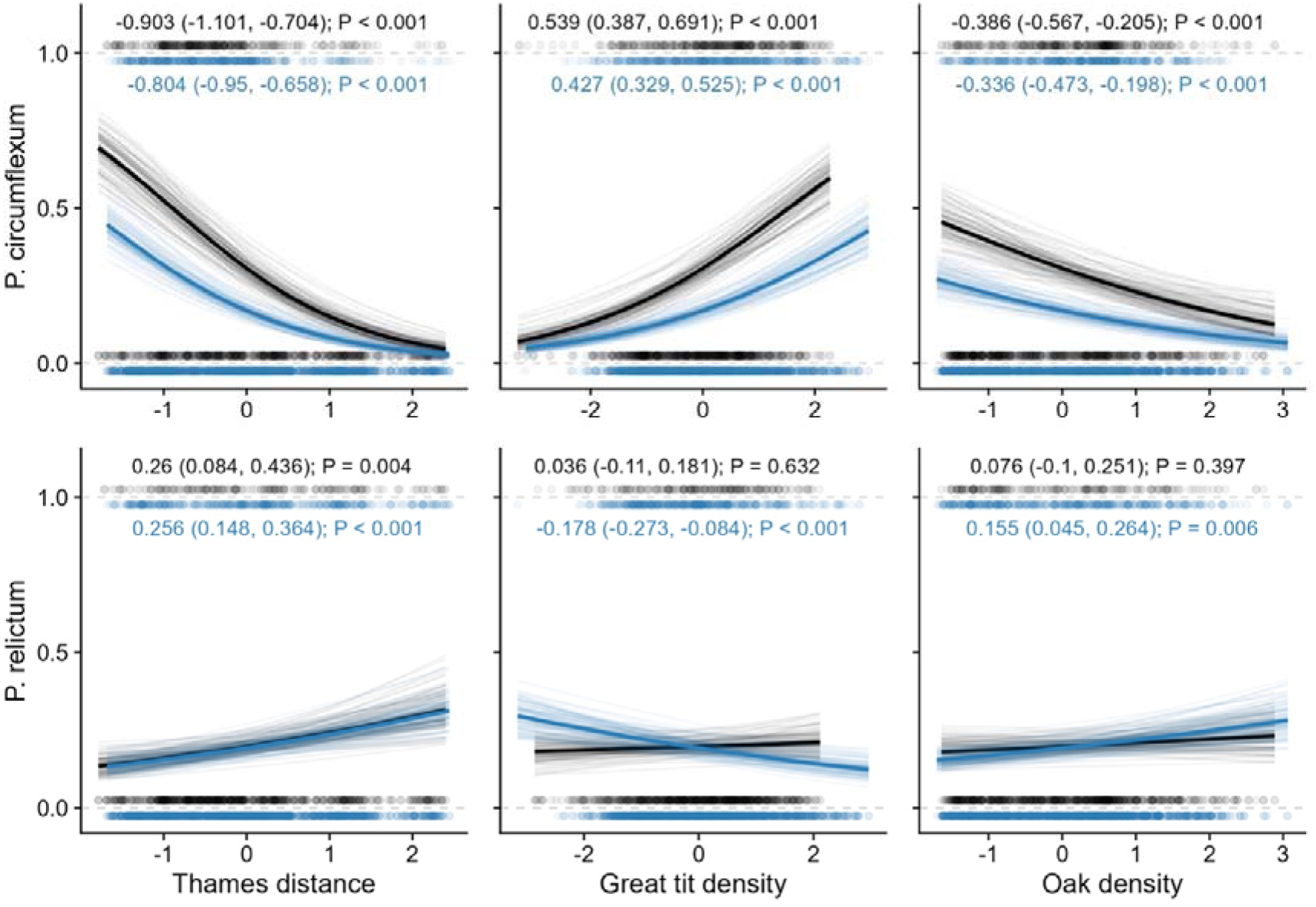
Modelled relationships between spatially distributed variables and avian malaria prevalence in great tits (black) and blue tits (blue). The dark line represents the mean of the posterior distribution for the model estimates; the light grey lines are 100 random draws from the posterior to represent uncertainty. Density refers to local annual density of great tits. Points denote observations of individual birds, with transparency to allow for visualisation of overplotting.

**Figure 5:**
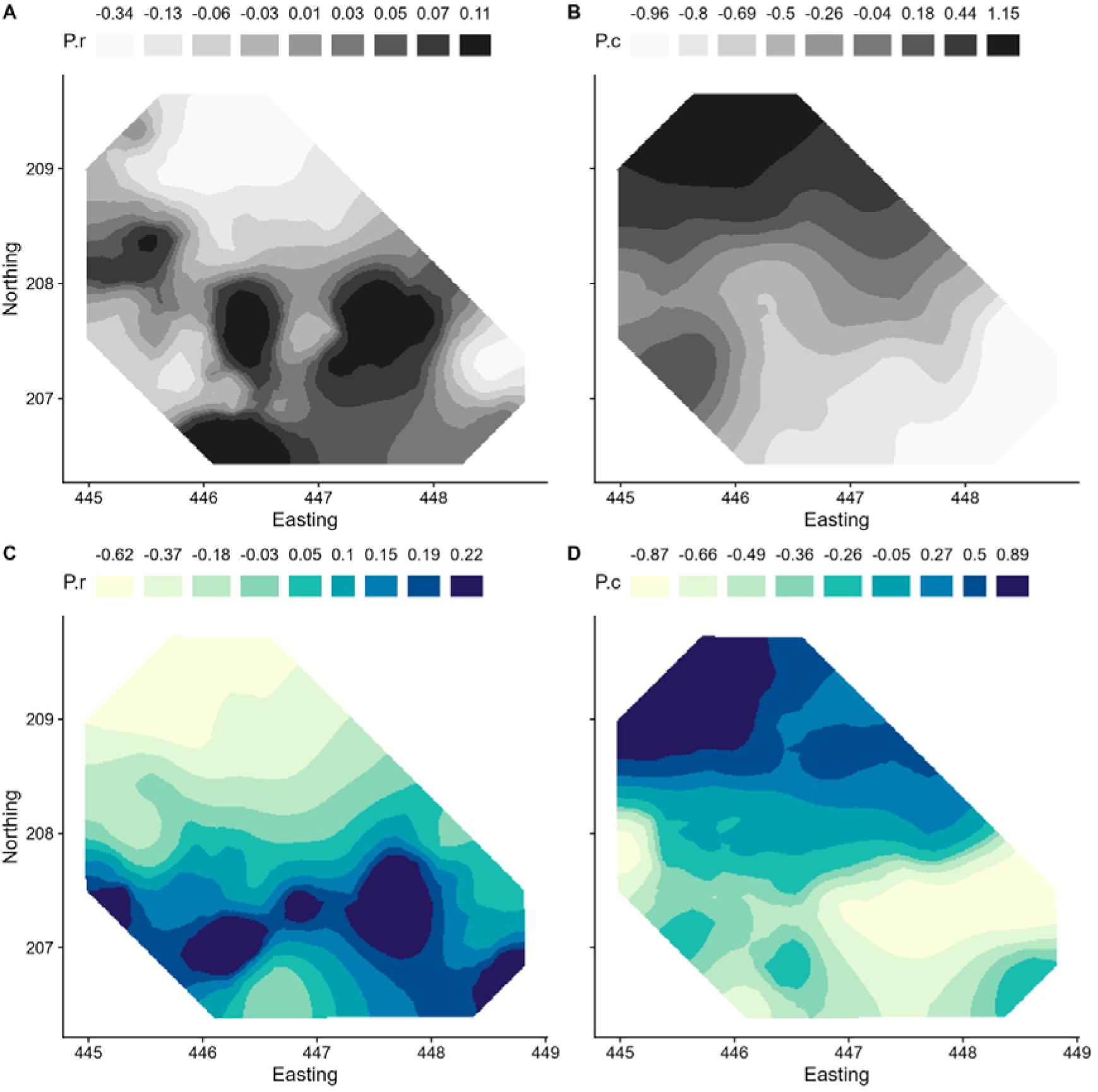
Projected spatial effects describing malaria prevalence in great tits (top row) and blue tits (bottom row). Shown are projections of the spatially-distributed SPDE random effect from the base model, without spatial fixed effects. Shading of the map denotes the lower bounds of nine quantiles of the spatial effects on the link scale, rounded to two decimal places, with darker colors representing higher malaria prevalence. Easting and Northing are in km.

While these effects likely represent the independent contributions of each variable after controlling for the others, several differences emerged between the full and single-variable models. Most notably, the association between oak density and P. circumflexum reversed direction in both host species. When fitted alone, oak density was positively associated with P. circumflexum prevalence in great tits (0.24 (0.12, 0.37), P < 0.001) and blue tits (0.17 (0.08, 0.26), P < 0.001), whereas in the full models it was negatively associated with prevalence in both great tits (–0.39 (–0.57, –0.20), P < 0.001) and blue tits (–0.34 (–0.47, –0.20), P < 0.001). Similarly, oak density was not associated with P. relictum prevalence in blue tits when fitted alone (–0.02 (–0.11, 0.07), P = 0.63), but became positively associated with prevalence in the full model (0.15 (0.04, 0.26), P = 0.01). In contrast, the effects of Thames distance and local annual great tit density were generally consistent between model types, although effect sizes were typically stronger in the full models.

Finally, models examining whether malaria infection explained brood failure found no evidence for an effect (ΔDIC < 2).

## Discussion

We uncovered species-specific associations between local density and brood failure in two wild bird species. In great tits, brood failure increased with local overall density of both or either species, whereas in blue tits brood failure decreased with local annual great tit density and annual combined density (but not with annual blue tit density). These differing associations suggest that brood failure’s density dependence may not be uniform even within the same community over the same timeframe, and likely reflect differences in how each species experiences local conditions. Beyond density, brood failure was strongly spatially structured: great tits showed higher failure farther from the Thames and in areas with lower oak density (with no evidence for comparable effects in blue tits). Finally, species-specific malaria prevalence was itself spatially and environmentally structured, with broadly similar patterns for *P. circumflexum* across host species and less consistent patterns for *P. relictum*, but parental infection status did not significantly explain brood failure in the subset of data with associated parasite screening. Taken together, these results suggest that complete brood failure reflects a spatially heterogeneous risk landscape that differs between sympatric species and is not straightforwardly explained by the measured habitat and disease variables alone.

Higher local density can impair reproductive performance through reduced per-capita resource availability, longer foraging distances, and intensified aggressive interactions among neighbours, all of which can reduce nestling growth and survival (Perrins 1965; Both 1998; Wilkin et al. 2006). When these pressures are further exacerbated by short-term constraints on resources, such as multi-day bad weather or temporary food shortfalls, provisioning may fall below a catastrophic viability threshold, leading to the sudden complete brood failures we observed. Lower brood failure in areas with higher oak density is consistent with the importance of resource availability, specifically oak-associated caterpillars in spring, with recent work further showing that caterpillar abundance and biomass are particularly high on oak and increase with oak foliage density within woodland stands (Macphie et al. 2025), and agrees with previously reported links between oak-rich territories, phenology, and breeding performance in this system and closely related ones (Perrins 1965; van Balen 1973; Morley et al. 2025; Jones et al. 2025). Parental depredation could also play a role: if high densities of great and blue tits attracts predators such as sparrowhawks, this could drive higher rates of parental loss, and brood failure events are consistent with such abrupt losses (Santema and Kempenaers 2018). Even without direct depredation, *perceived* greater predation risk by adults could also alter time allocation, reduce provisioning, and shift foraging decisions (Lima 2009; McMahon et al. 2024), ultimately reducing offspring production (Zanette et al. 2011). Further, density dependence can arise when individuals compete for limiting scarce refuges from predators, creating spatially structured survival outcomes (Hurley, Hebblewhite, and Gaillard 2020). However, higher prey density can also *dilute* risk, adding complexity. For example, per-capita sparrowhawk predation on great tits likely decreases with increasing breeding density in other populations (Götmark and Andersson 2005). Such complexities could explain our discovery of both positive and negative effects of local density on brood failure. Although our data rarely uncovered predation events on nestlings, this does not rule out predator-mediated pathways operating through adult mortality or risk-induced reductions in provisioning (Lima 2009; Zanette et al. 2011; Voelkl et al. 2016; Santema and Kempenaers 2018), and further work could now attempt to directly identify these pathways.

Beyond species-specific insight into brood failure mechanisms, a key finding is that local density’s effects can differ even between sympatric, closely related, and interacting species in the same community. Specifically, while brood failure increased with local overall density in great tits, it declined in blue tits with local annual great tit and combined density, consistent with the possibility that annual density in blue tits reflects favourable within-year conditions rather than reflecting crowding-driven increases in catastrophic failure. For instance, if annual density partly reflects settlement into locally attractive areas before breeding, because birds use early-season cues or because some patches consistently provide better resources, then denser neighbourhoods could also be those with lower failure risk, producing a negative association between annual density and complete failure even when competition is present. This is particularly plausible given that the negative association is observed for local annual great tit density (or annual combined density) rather than annual blue tit density, suggesting that great tit density may be acting partly as an index of local habitat quality or settlement attractiveness rather than simply as a measure of competitive pressure. If local annual great tit density tracks within-year habitat quality or resource pulses more closely, it may better reflect conditions experienced by blue tits than blue tit density itself. Additionally, blue tits might experience negative crowding/competition effects mainly through partial reductions (e.g. smaller broods, slower growth, fewer fledglings) rather than increased risk of complete brood failure. Under this scenario, total failure may be lowered where annual density is higher if such sites represent the best within-year conditions, and then negative effects of density dependence may be expressed primarily in conditional outcomes among surviving broods. Annual density can also covary with mating dynamics in blue tits (including extra-pair paternity), which may affect parental investment, brood size, and failure risk, though evidence for density effects is mixed across studies (Charmantier and Perret 2004; Mennerat et al. 2018). A potential next step in this sense could therefore be to examine the fine-scale mechanisms whereby blue tits may show negative density dependence in fledgling number conditional on fledging at least one chick, even if complete failure declines with annual density.

The negative association between local annual great tit density and blue tit brood failure could reflect dominance-mediated sorting, given what is known about their interspecific interactions. Great tits can be strong interference competitors for nest sites (due to their larger size), while blue tits can be effective exploitative competitors during breeding, particularly where nest sites are not limiting (Minot 1981; Minot and Perrins 1986; Dhondt 2010). If high local annual great tit density is concentrated in favourable/attractive parts of the woodland, the blue tits that breed there may be a non-random subset (either higher-quality individuals, or individuals occupying nest sites that are less prone to total brood failures). Conversely, if interspecific interference excludes the lowest-quality blue tits from high-density neighbourhoods, this could reduce the fraction of blue tit attempts that are vulnerable to complete failure even if competition still reduces other components of performance. Interspecific social information transfer could also contribute: if birds can take cues from heterospecifics, this may influence space use and risk-sensitive behaviour (Magrath et al. 2015; Firth et al. 2016; Keen et al. 2020), and local population size can also reshape social structure in ways that plausibly alter information landscapes (Beck et al. 2023; Beck et al. 2024). If blue tits benefit disproportionately from such information in high great tit density neighbourhoods, local annual great tit density could be associated with reduced total failure without implying improved resource conditions per se. Finally, predators could likewise mediate this effect as referenced above: for example, if great tits are easier prey to sparrowhawks (again due to their size), areas of high great tit density might in fact specifically dilute the risk of predation for blue tits, reducing their risk and therefore reducing rates of brood failure.

Despite showing strong spatial heterogeneity and clear associations with local density and habitat, species-specific malaria infection status did not explain brood failure in either host species. This was the case even though *P. circumflexum* and *P. relictum* showed distinct ecological patterns, and even though several infection models were improved by accounting for residual spatial autocorrelation. It is possible that malaria mainly imposes costs on other fitness components, acting too gradually to lead to total brood failure (Lachish et al. 2011). Alternatively, binary infection status may remain too coarse even when parasite species are separated: costs can depend on infection intensity, infection stage, and timing relative to breeding, and chronic low-intensity infections may have limited direct impact on acute nestling survival (Atkinson and van Riper 1991; Knowles et al. 2011; Knowles et al. 2014). As such, malaria remains informative as a spatially structured marker of exposure processes and habitat gradients, and may covary with the same environmental gradients that shape breeding density and failure risk. Although avian malaria can reduce reproductive success in some bird systems (Marzal et al. 2005; Lachish et al. 2011), we do not find evidence that parental infection status is a primary driver of complete brood collapse, or that it explains the observed effects of local density on brood failure.

Finally, we uncovered substantial further spatial variation in brood failure beyond population and oak density. Higher failure farther from the Thames suggests broad spatial gradients in other factors such as microclimate, habitat structure, and/or prey availability may exist which are not fully captured by our included habitat-based covariates. We also uncovered substantial spatial heterogeneity in the models; in wild populations such as great tits and blue tits, such residual structure (Gokcekus et al. 2023; Woodman et al. 2026) could likewise indicate unmeasured habitat features such as fine-scale phenology (Jones et al. 2025), canopy structure, microclimate, or spatial variation in unobserved predator activity or ectoparasite communities (known to shape among-site differences in reproductive rate in other bird systems (e.g. Grant et al. 1999; Wilson and Arcese 2006)), that are only partly correlated with our covariates. That spatial structure remains strong after including local density, habitat proxies, and species-specific infection statuses suggests that density effects operate within a broader, spatially heterogeneous landscape of baseline risks of complete brood failure. While more work is needed to draw definitive conclusions about how these and other habitat features act as direct causal drivers, the combination of spatial effects is consistent with great tit failure being shaped by spatially structured resources and with microclimatic constraints, with density-related competition increasing the probability of threshold-type collapse (i.e. total brood failure) on top of the expected baseline.

Together, our results provide a foundation for future work aimed at more direct mechanistic investigations to uncover fine-scale drivers of complete brood failure. For instance, to further examine the relative importance of resource limitation and phenological mismatch, integrating measures of local spring phenology (Cole et al. 2021) and caterpillar availability could test whether density effects increase in mismatched years or in habitats with lower resource availability. Similarly, testing interactions between density and habitat proxies (oak density, distance to the Thames) could probe further into the resource-limitation interpretation for great tits, for example by testing whether density dependence is strongest where resources are most limiting or microclimate is most challenging.

Additionally, further mechanistic analyses could include demographic characteristics as additional interacting variables to examine whether density effects are stronger in young females, late broods, or years with poorer phenological matching. On top of this, if independent spatial map layers were collected and included (e.g., habitat maps, phenology indices, predator monitoring), comparing these with the currently fitted spatial failure surface could also help identify which unmeasured gradients may be the most plausible candidates for the strong residual spatial structure. Finally, experimental approaches may also be possible for manipulating local density, such as through direct nestbox density manipulations (Both 1998) or by altering the density of individuals during the non-breeding season (Beck et al. 2024) which may be expected to then carry-over into the subsequent breeding positions (Firth & Sheldon 2016), which could help definitively separate crowding/competitive effects from settlement choices/bias.

Beyond its implications for ecological applications such as monitoring and demographic forecasting, brood failure also has implications for wild animal welfare because it entails the death of multiple nestlings, and all plausible proximate causes (starvation, exposure, or prolonged stress following reduced provisioning) are likely to involve substantial suffering prior to death. Physiological work across taxa shows that starvation and repeated food limitation trigger strong metabolic and endocrine responses, and that impacts can extend beyond the immediate deprivation period (McCue et al. 2017). As such, by identifying drivers and indicators of brood failure rates, our results help inform the distribution of welfare-relevant events in the wild.

Overall, our results show that complete brood failure is a spatially structured outcome, of which the ecological correlates can differ even between closely related species co-occurring in the same environment. In great tits, failure risk was highest with dense local populations and in areas associated with lower resource conditions, consistent with local constraints increasing the likelihood of brood collapse. In blue tits, failure declined with local annual great tit and combined density, suggesting that within-year habitat quality, settlement processes, and interspecific context can shape complete brood failure in ways that do not line up to simple intraspecific crowding costs. More broadly, these findings show that density dependence in complete brood failure is species– and context-dependent, and suggest that understanding when complete brood failure occurs requires information on environmental factors, community composition and individual distributions, and residual spatial structure.

## Acknowledgements

GFA acknowledges funding from NSF grant DEB-2211287 and WAI (C-2023-00057). JAF acknowledges funding from BBSRC (BB/S009752/1), NERC (NE/S010335/1 and NE/V013483/2) and WAI (C-2023-00057). The long-term population studies in Wytham have been supported by numerous funding sources, including recently by grants from BBSRC (BB/L006081/1), ERC (AdG 250164), NERC (NE/K006274/1, NE/S010335/1 and NE/X000184/1) and a UKRI Frontiers Award (EP/X024520/1). Malaria diagnoses were funded by NERC grant NE/F005725/1.

## Supplementary Information

**Table S1.** Measures of parental malaria infection status used in the analyses. Rows show the host species, years for which malaria screening data were available, and the parasite-specific infection measure analysed. All measures were treated as binary infection variables, with individuals coded as infected or uninfected for the relevant *Plasmodium* species. Malaria diagnoses were based on PCR-based screening of blood samples from breeding adults sampled during the nestling period. Great tit samples were available for 2008–2009 and blue tit samples for 2005–2010. Infections were assigned to *P. relictum* or *P. circumflexum*, the two dominant *Plasmodium* morphospecies in this system, using diagnostic information from qPCR product melting temperature and/or *cytochrome b* sequence data, depending on the assay used. The qPCR assay targeted a 188-bp fragment of *Plasmodium cytochrome b* and was run in triplicate, with samples scored positive if any replicate amplified; some blue tit diagnoses also used nested *cytochrome b* PCR following Waldenström et al. (2004). Where parental infection status was used in brood-failure models, nests were included only when malaria data were available for at least one parent. References for the diagnostic methods and previous use of these data in the Wytham tit system are given in the main text.

| Host species | Years | Measure analysed | N | Prop. Infected |
| --- | --- | --- | --- | --- |
| Great tit | 2008-2009 | *P. relictum* | 1146 | 20.41% |
| Great tit | 2008-2009 | *P. circumflexum* | 1146 | 34.03% |
| Blue tit | 2005-2010 | *P. relictum* | 2994 | 20.44% |
| Blue tit | 2005-2010 | *P. circumflexum* | 2994 | 19.97% |

**Figure S1.**
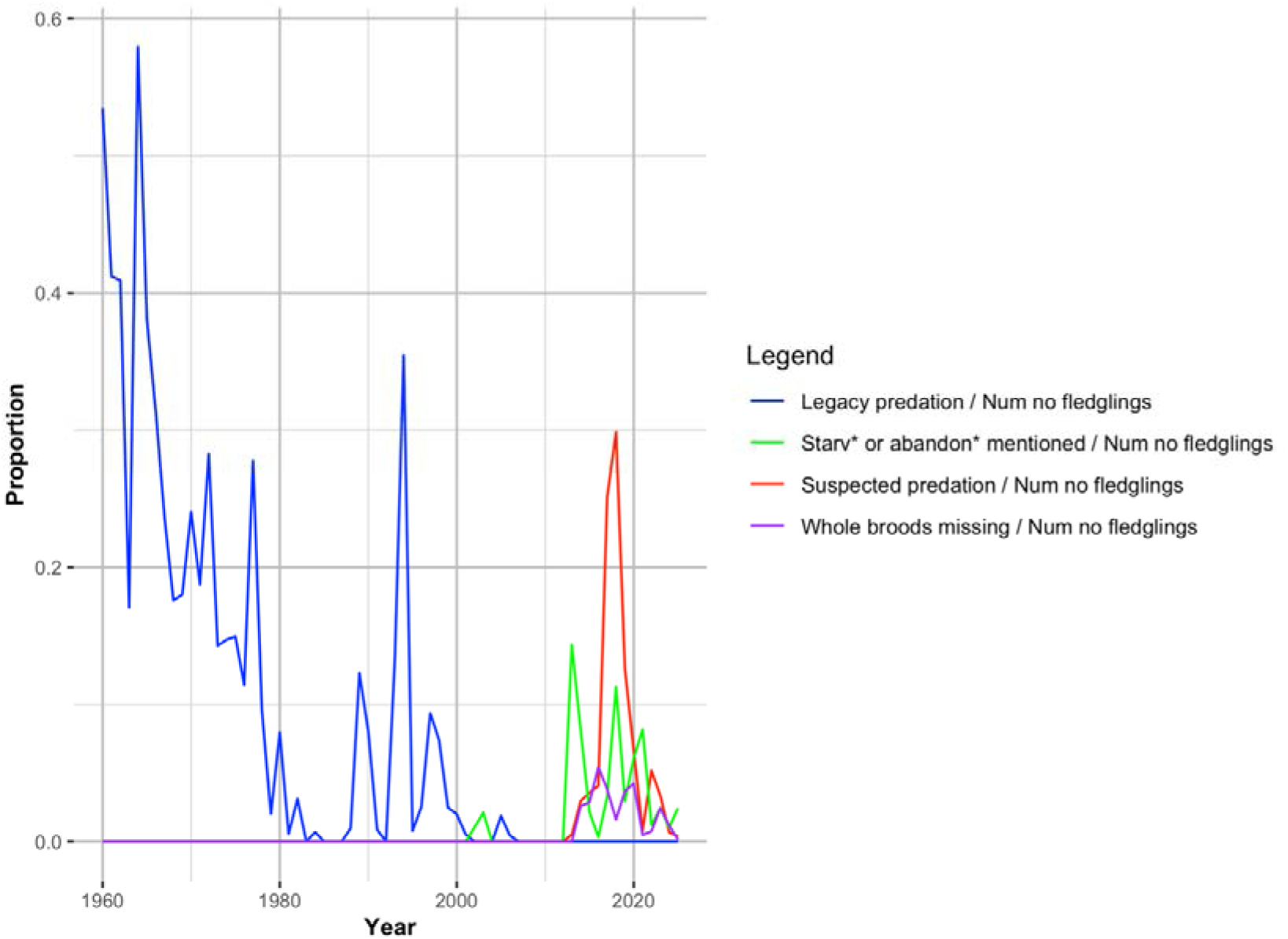
Annual proportions of 0-fledgling breeding attempts with specific failure annotations. For each year (both species combined), lines show the proportion of 0-fledgling breeding attempts that carried each annotation, calculated as (annual count with annotation)/(annual count with 0 fledglings): legacy predation (blue), suspected predation (red), whole broods missing (purple), and comments containing “starv*” or “abandon*” (green). Proportions can vary strongly in years with low numbers of 0-fledgling attempts and should be interpreted alongside the corresponding annual counts (see Fig. S2).

**Figure S2.**
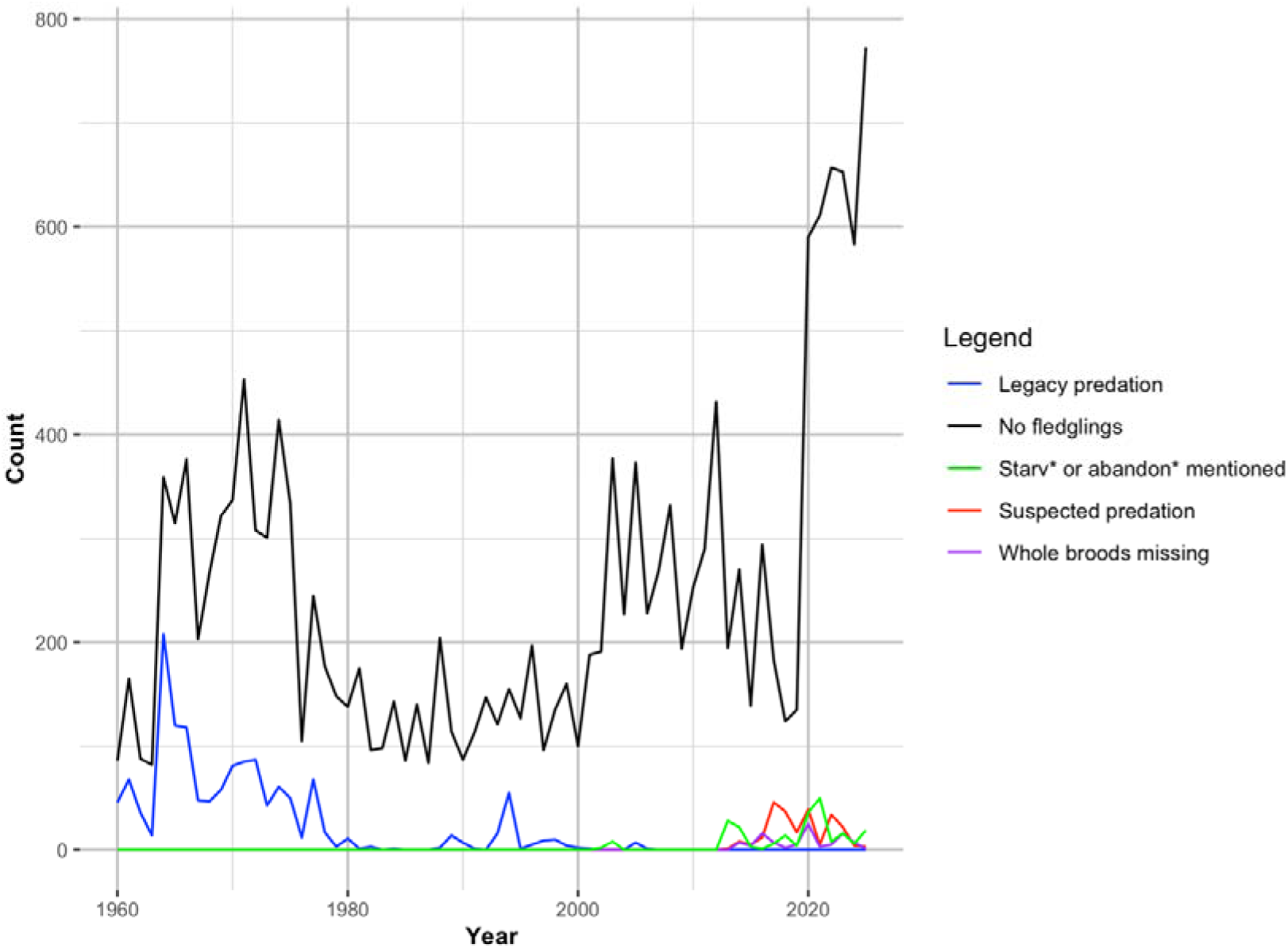
Annual counts of failure-related annotations among 0-fledgling breeding attempts. For each year (both species combined), the black line shows the number of breeding attempts with 0 fledglings recorded. Coloured lines show, among those 0-fledgling attempts, the annual count carrying each annotation: “legacy predation” flag (blue), “suspected predation” flag (red), “whole broods missing” flag (purple), and free-text comments containing “starv*” or “abandon*” (green).

**Figure S3.**
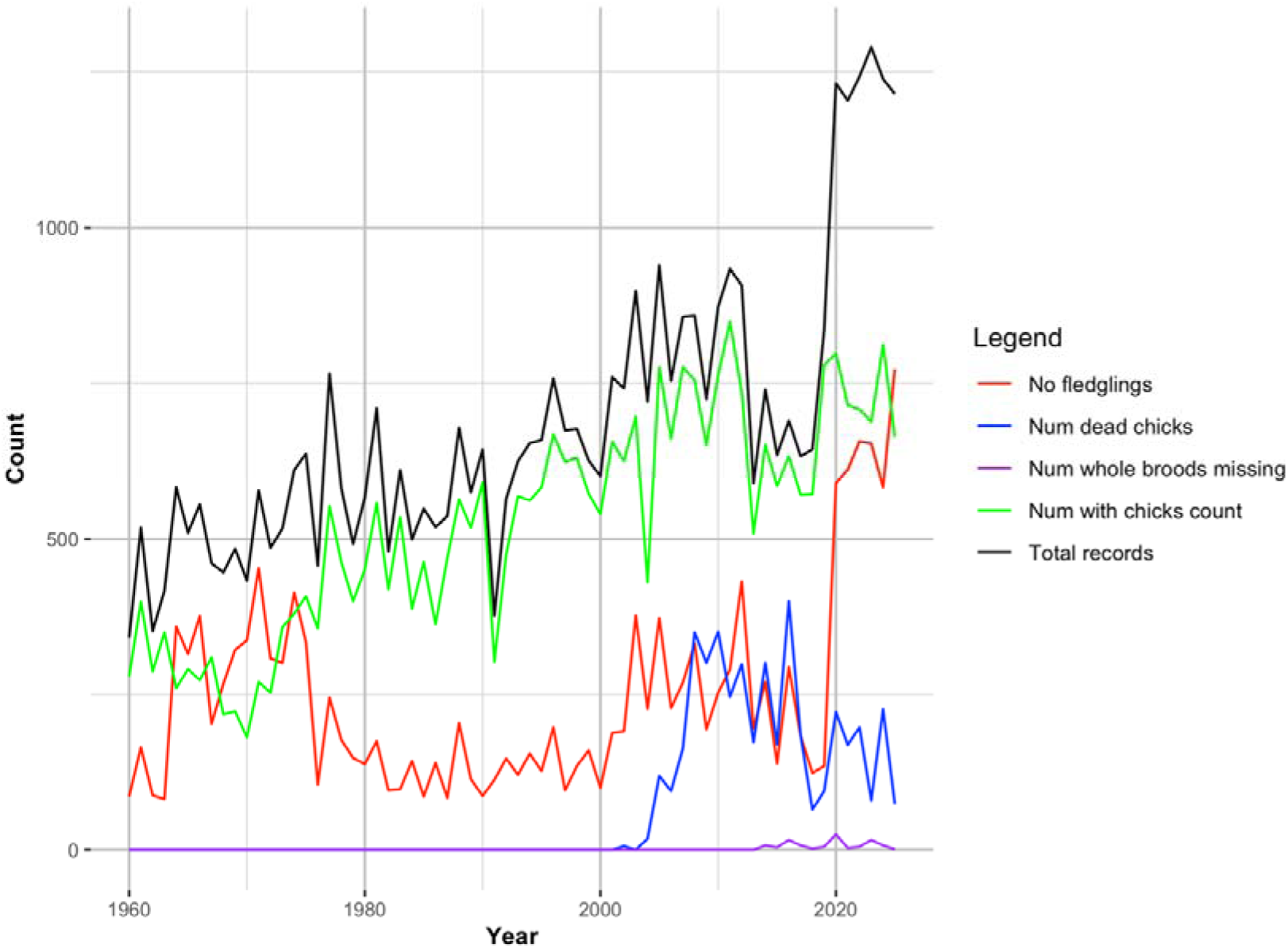
Annual summary of brood-record annotations used to contextualise 0-fledgling outcomes. Time series for great tits and blue tits combined showing: total number of breeding-attempt records (“Total records”, black); number of attempts with 0 fledglings recorded (“No fledglings”, red); number of attempts with >0 dead chicks recorded in the nest (“Num dead chicks”, blue); number of attempts flagged as “whole broods missing” (“Num whole broods missing”, purple); and number of attempts with a chick count recorded at any point (“Num with chicks count”, green).

## Notes

### Competing Interest Statement

The authors have declared no competing interest.

## References

1. Albery, Gregory F. 2022. “Density Dependence and Disease Dynamics: Moving towards a Predictive Framework.” EcoEvoRxiv Preprints. 10.32942/OSF.IO/GAW49

2. Albery, Gregory F., Chris Newman, Julius Bright Ross, David W. Macdonald, Shweta Bansal, and Christina D. Buesching. 2020. “Negative Density-Dependent Parasitism in a Group-Living Carnivore.” Proceedings of the Royal Society B: Biological Sciences 287 (1941): 20202655.

3. Albery, G., Becker, D., Firth, J. A., et al. 2025. “Density-dependent network structuring within and across wild animal systems.” Nature Ecology & Evolution. 10.1038/s41559-025-02843-z

4. Atkinson, Carter T., and Charles van Riper III. 1991. “Pathogenicity and Epizootiology of Avian Haematozoa: Plasmodium, Leucocytozoon, and Haemoproteus.” In Bird-Parasite Interactions: Ecology, Evolution, and Behaviour, edited by J. E. Loye and M. Zuk, 19–48. Oxford, UK: Oxford University Press.

5. Beck, K., Farine, D., Firth, J. A., and Sheldon, B. 2023. “Variation in local population size predicts social network structure in wild songbirds.” Journal of Animal Ecology. 10.1111/1365-2656.14015

6. Beck, K., Regan, C., McMahon, K., Crofts, S., Cole, E., Firth, J. A., and Sheldon, B. 2024. “Experimental manipulation of local population density in wild great tits (Parus major) alters local social structure but has little effect on information acquisition.” Animal Behaviour. 10.1016/j.anbehav.2023.12.010

7. Both, Christiaan. 1998. “Experimental Evidence for Density Dependence of Reproduction in Great Tits.” Journal of Animal Ecology 67 (4): 667–674. 10.1046/j.1365-2656.1998.00228.x

8. Byholm, P. 2005. “Site-Specific Variation in Partial Brood Loss in Northern Goshawks.” Annales Zoologici Fennici 42: 81–90.

9. Charmantier, Anne, and Philippe Perret. 2004. “Manipulation of Nest-box Density Affects Extra-pair Paternity in a Population of Blue Tits (Parus caeruleus).” Behavioral Ecology and Sociobiology 56: 360–365. 10.1007/s00265-004-0794-5

10. Charnov, Eric L., and John R. Krebs. 1974. “On Clutch-size and Fitness.” Ibis 116 (2): 217–219.

11. Clark, Anne Barrett, and David Sloan Wilson. 1981. “Avian Breeding Adaptations: Hatching Asynchrony, Brood Reduction, and Brood Failure.” The Quarterly Review of Biology 56 (3): 253–277.

12. Cody, Martin L. 1966. “A General Theory of Clutch Size.” Evolution 20 (2): 174–184.

13. Cole, E.F., Regan, C.E. & Sheldon, B.C. Spatial variation in avian phenological response to climate change linked to tree health. Nature Climate Change. 11, 872–878 (2021). 10.1038/s41558-021-01140-4

14. Dhondt, André A. 2010. “Effects of Competition on Great and Blue Tit Reproduction: Intensity and Importance in Relation to Habitat Quality.” Journal of Animal Ecology 79 (1): 257–265. 10.1111/j.1365-2656.2009.01624.x

15. Firth, J. A., et al. 2016. “Pathways of information transmission amongst wild songbirds follow experimentally imposed changes to social foraging structure.” Biology Letters 12: 20160144. 10.1098/rsbl.2016.0144

16. Firth, J. A., et al. 2018. “Spatial, temporal and demographic based differences in wild great tits’ nest-site visits and the consequences for reproduction.” Journal of Avian Biology. 10.1111/jav.01740

17. Firth, J. A., and Sheldon, B. C. 2016. “Social carry-over effects underpin trans-seasonally linked structure in a wild bird population.” Ecology Letters 19: 1324–1332. 10.1111/ele.12669

18. Gamelon, M., Firth, J. A., Le Moullec M., Petry W.K., and Ssalguero-Gomez, R. 2021. “Longitudinal demographic data collection.” In Demographic Methods across the Tree of Life. Oxford University Press. 10.1093/oso/9780198838609.003.0005

19. Gokcekus S, Firth JA et al. 2025. Different types of social links contrastingly shape reproductive traits in a multi-level society of wild songbirds. Behavioural Ecology and Sociobiology; DOI: 10.1007/s00265-025-03594-4

20. Gokcekus S, Firth JA et al. 2023. Social familiarity and spatially variable environments independently determine reproductive fitness in a wild bird. American Naturalist; DOI: 10.1086/724382

21. Gosler, Andrew. 1993. The Great Tit. Hamlyn Species Guides. London, England: Hamlyn.

22. Götmark, Frank, and Malte Andersson. 2005. “Predation by Sparrowhawks Decreases with Increased Breeding Density in a Songbird, the Great Tit.” Oecologia 142 (2): 177–183. 10.1007/s00442-004-1715-z

23. Grant, Murray C., Chris Orsman, Jon Easton, Chris Lodge, Malcolm Smith, Guy Thompson, Stephen Rodwell, and Niall Moore. 1999. “Breeding Success and Causes of Breeding Failure of Curlew Numenius Arquata in Northern Ireland.” Journal of Applied Ecology 36 (1): 59–74.

24. Hasik, Adam Z., Shane Butt, Richard S. Turner, Katie Maris, Sean Morris, Ali Morris, Josephine M. Pemberton, and Gregory F. Albery. 2024. “Population Density Drives Increased Parasitism via Greater Exposure and Reduced Resource Availability in Wild Hosts.” bioRxiv. 10.1101/2024.07.08.602460

25. Hurley, Mark A., Mark Hebblewhite, and Jean-Michel Gaillard. 2020. “Competition for Safe Real Estate, Not Food, Drives Density-Dependent Juvenile Survival in a Large Herbivore.” Ecology and Evolution 10 (12): 5464–5475.

26. Hurtrez-Boussès, Sylvie, Philippe Perret, François Renaud, and Jacques Blondel. 1997. “High Blowfly Parasitic Loads Affect Breeding Success in a Mediterranean Population of Blue Tits.” Oecologia 112 (4): 514–517.

27. Jones, C. V., Regan, C., Cole, E. F., Firth, J. A., and Sheldon, B. C. 2025. “Shared environmental similarity between relatives influences heritability of reproductive timing in wild great tits.” Evolution. 10.1093/evolut/qpae155

28. Jones, C. V. (2025). Spatial, social and environmental drivers of the timing of reproduction in the wild [DPhil thesis, University of Oxford]. Oxford University Research Archive. DOI: 10.5287/ora-8jozbjg2o.

29. Julliard, Romain, Robin H. McCleery, Jean Clobert, and Christopher M. Perrins. 1997. “Phenotypic Adjustment of Clutch Size Due to Nest Predation in the Great Tit.” Ecology 78 (2): 394–404. 10.1890/0012-9658(1997)078[0394:PAOCSD]2.0.CO;2

30. Karaer, Mina Cansu, Nina Čebulj-Kadunc, and Tomaž Snoj. 2023. “Stress in Wildlife: Comparison of the Stress Response among Domestic, Captive, and Free-Ranging Animals.” Frontiers in Veterinary Science 10: 1167016.

31. Keen, Sara C., Ella F. Cole, Michael J. Sheehan, and Ben C. Sheldon. 2020. “Social Learning of Acoustic Anti-Predator Cues Occurs between Wild Bird Species.” Proceedings of the Royal Society B: Biological Sciences 287 (1920): 20192513. 10.1098/rspb.2019.2513

32. Kirkpatrick, Chris, C. Conway, and M. H. Ali. 2009. “Sanitation of Entire Broods of Dead Nestlings May Bias Cause-Specific Brood Failure Rates.” Ibis 151: 207–211.

33. Kochert, Michael N., Karen Steenhof, and Jessi L. Brown. 2019. “Effects of Nest Exposure and Spring Temperatures on Golden Eagle Brood Survival: An Opportunity for Mitigation.” Journal of Raptor Research 53(1):91–97 10.3356/JRR-17-100

34. Knowles, Sarah C. L., Matthew J. Wood, Ricardo Alves, Teddy A. Wilkin, Staffan Bensch, and Ben C. Sheldon. 2011. “Molecular Epidemiology of Malaria Prevalence and Parasitaemia in a Wild Bird Population.” Molecular Ecology 20 (5): 1062–1076. 10.1111/j.1365-294X.2010.04909.x

35. Knowles, Sarah C. L., Matthew J. Wood, Frédéric A. B. V. Alves, and Ben C. Sheldon. 2014. “Dispersal in a Patchy Landscape Reveals Contrasting Determinants of Infection in a Wild Avian Malaria System.” Journal of Animal Ecology 83 (2): 429–439. 10.1111/1365-2656.12154

36. Lachish, Shelly, Sarah C. L. Knowles, Frédéric A. B. V. Alves, Matthew J. Wood, and Ben C. Sheldon. 2011. “Fitness Effects of Endemic Malaria Infections in a Wild Bird Population: The Importance of Ecological Structure.” Journal of Animal Ecology 80 (6): 1196–1206. 10.1111/j.1365-2656.2011.01836.x

37. Lachish, Shelly, Sarah C. L. Knowles, Ricardo Alves, Irem Sepil, Alicia Davies, Simon Lee, Matthew J. Wood, and Ben C. Sheldon. 2013. “Spatial Determinants of Infection Risk in a Multi-Species Avian Malaria System.” Ecography 36 (5): 587–598. 10.1111/j.1600-0587.2012.07811.x.

38. Lima, Steven L. 2009. “Predators and the Breeding Bird: Behavioral and Reproductive Flexibility under the Risk of Predation.” Biological Reviews 84 (3): 485–513. 10.1111/j.1469-185X.2009.00085.x

39. Lindgren, Finn, Håvard Rue, and Johan Lindstrom. 2011. “An Explicit Link between Gaussian Fields and Gaussian Markov Random Fields: The Stochastic Partial Differential Equation Approach.” Journal of the Royal Statistical Society: Series B (Statistical Methodology) 73 (4): 423–498.

40. Lindgren, Finn, and Håvard Rue. 2015. “Bayesian Spatial Modelling with R-INLA.” Journal of Statistical Software 63: 1–25.

41. Macphie, Kirsty H., Jelmer M. Samplonius, Jarrod D. Hadfield, James W. Pearce-Higgins, and Albert B. Phillimore. 2025. “Tree Taxon Effects on the Phenology of Caterpillar Abundance and Biomass.” Oikos 2025 (4): e10972. 10.1111/oik.10972

42. Magrath, Robert D., Tonya M. Haff, Pamela M. Fallow, and Andrew N. Radford. 2015. “Eavesdropping on Heterospecific Alarm Calls: From Mechanisms to Consequences.” Biological Reviews 90 (2): 560–586. 10.1111/brv.12122

43. Marzal, Alfonso, Florentino de Lope, Carlos Navarro, and Anders Pape Møller. 2005. “Malarial Parasites Decrease Reproductive Success: An Experimental Study in a Passerine Bird.” Oecologia 142 (4): 541–545. 10.1007/s00442-004-1757-2

44. Martin, Thomas E. 1995. “Avian Life History Evolution in Relation to Nest Sites, Nest Predation, and Food.” Ecological Monographs 65 (1): 101–127.

45. Matthysen, Erik, and Frank Adriaensen. 1998. “Forest Size and Isolation Have No Effect on Reproductive Success of Eurasian Nuthatches (Sitta Europaea).” The Auk 115 (4): 955–963.

46. McCue, Marshall D., John S. Terblanche, and Joshua B. Benoit. 2017. “Learning to Starve: Impacts of Food Limitation beyond the Stress Period.” The Journal of Experimental Biology 220 (Pt 23): 4330–4338.

47. McMahon, K., et al. 2024. “Social network centrality predicts dietary decisions in a wild bird population.” iScience. 10.1016/j.isci.2024.109581

48. Mennerat, Adèle, Anne Charmantier, Christian Jørgensen, and Sigrunn Eliassen. 2018. “Correlates of Complete Brood Failure in Blue Tits: Could Extra-pair Mating Provide Unexplored Benefits to Females?” Journal of Avian Biology 49 (5). 10.1111/jav.01701

49. Milchev, B., and V. Georgiev. 2012. “Plastic Fibres Cause a Brood Failure in a Long-Legged Buzzard Buteo Rufinus Nest.” Acrocephalus. https://ptice.si/wp-content/uploads/2014/04/acrocephalus_32_150-151.pdf#page=97

50. Minot, Edward O. 1981. “Effects of Interspecific Competition for Food in Breeding Blue and Great Tits.” Journal of Animal Ecology 50 (2): 375–385. 10.2307/4061

51. Minot, Edward O., and C. M. Perrins. 1986. “Interspecific Interference Competition—Nest Sites for Blue and Great Tits.” Journal of Animal Ecology 55 (1): 331–350. 10.2307/4712

52. Morley, Lucy M., Sam J. Crofts, Ella F. Cole, and Ben C. Sheldon. 2025. “Quantifying Phenology in the Deciduous Tree and Phytophagous Insect System: A Methodological Comparison.” Ecology and Evolution 15 (9): e71821. 10.1002/ece3.71821

53. Mudge, G. P., and T. R. Talbot. 1993. “The Breeding Biology and Causes of Brood Failure of Scottish Black-throated Divers Gavia arctica.” Ibis 135 (2): 113–120. 10.1111/j.1474-919X.1993.tb02822.x

54. Palinauskas, Vaidas, Vlad Kosarev, Anatoly Shapoval, Staffan Bensch, and Gediminas Valkinas. 2007. “Comparison of Mitochondrial Cytochrome B Lineages and Morphospecies of Two Avian Malaria Parasites.” Zootaxa 1626 (1): 39–50.

55. Payne, R. B. 1977. “The Ecology of Brood Parasitism in Birds.” Annual Review of Ecology and Systematics 8: 1–28. 10.1146/annurev.es.08.110177.000245

56. Perrins, C. M. 1965. “Population Fluctuations and Clutch-size in the Great Tit, Parus major L.” Journal of Animal Ecology 34 (3): 601–647. 10.2307/2453

57. Rue, Håvard, Sara Martino, and Nicolas Chopin. 2009. “Approximate Bayesian Inference for Latent Gaussian Models by Using Integrated Nested Laplace Approximations.” Journal of the Royal Statistical Society: Series B (Statistical Methodology) 71 (2): 319–392. 10.1111/j.1467-9868.2008.00700.x

58. Santema, Peter, and Bart Kempenaers. 2018. “Complete Brood Failure in an Altricial Bird Is Almost Always Associated with the Sudden and Permanent Disappearance of a Parent.” Journal of Animal Ecology 87 (5): 1239–1250.

59. Schöll, Eva Maria, and Sabine Marlene Hille. 2020. “Heavy and Persistent Rainfall Leads to Brood Reduction and Brood Failure in a Passerine Bird.” Journal of Avian Biology 51 (7). 10.1111/jav.02418

60. Sih, Andrew. 1984. “Optimal Behavior and Density-Dependent Predation.” The American Naturalist 123 (3): 314–326.

61. Slagsvold, Tore. 1981. “Clutch Size and Population Stability in Birds: A Test of Hypotheses.” Oecologia 49 (2): 213–217.

62. Stearns, Stephen C. 1992. The Evolution of Life Histories. London, England: Oxford University Press.

63. Sutherland, William J. 1996. From Individual Behaviour to Population Ecology. Oxford, UK: Oxford University Press.

64. Thibault, Janet M., Felicia J. Sanders, and Patrick G. R. Jodice. 2010. “Parental Attendance and Brood Success in American Oystercatchers in South Carolina.” Waterbirds 33 (4): 511–517.

65. van Balen, J. H. 1973. “A Comparative Study of the Breeding Ecology of the Great Tit Parus major in Different Habitats.” Ardea 61 (1-2): 1–93. 10.5253/arde.v61.p1

66. Voelkl, B., Firth, J. A., and Sheldon, B. C. 2016. “The socio-ecology of fear: nonlethal predator effects on the social composition of wild bird flocks.” Scientific Reports 6: 33476. 10.1038/srep33476

67. Waldenström, Jonas, Staffan Bensch, Dennis Hasselquist, and Örjan Östman. 2004. “A New Nested Polymerase Chain Reaction Method Very Efficient in Detecting Plasmodium and Haemoproteus Infections from Avian Blood.” Journal of Parasitology 90 (1): 191–194. 10.1645/GE-3221RN

68. Wang, Guiming, N. Thompson Hobbs, Saran Twombly, Randall B. Boone, Andrew W. Illius, Iain J. Gordon, and John E. Gross. 2009. “Density Dependence in Northern Ungulates: Interactions with Predation and Resources.” Population Ecology 51 (1): 123–132.

69. Wilkin, Teddy A., Dany Garant, Andrew G. Gosler, and Ben C. Sheldon. 2006. “Density Effects on Life-History Traits in a Wild Population of the Great Tit Parus major: Analyses of Long-Term Data with GIS Techniques.” Journal of Animal Ecology 75 (2): 604–615. 10.1111/j.1365-2656.2006.01078.x

70. Williams, George C. 1966. “Natural Selection, the Costs of Reproduction, and a Refinement of Lack’s Principle.” The American Naturalist 100 (916): 687–690.

71. Wilson, Scott, and Peter Arcese. 2006. “Nest Depredation, Brood Parasitism, and Reproductive Variation in Island Populations of Song Sparrows (Melospiza Melodia).” The Auk 123 (3): 784–794.

72. Wood, Matthew J., Catherine L. Cosgrove, Teddy A. Wilkin, Sarah C. L. Knowles, Karen P. Day, and Ben C. Sheldon. 2007. “Within-Population Variation in Prevalence and Lineage Distribution of Avian Malaria in Blue Tits, Cyanistes Caeruleus.” Molecular Ecology 16 (15): 3263–3273.

73. Woodman, J., Firth, J. A., Cole, E., and Sheldon, B. C. 2026. “Age-specificity in territory quality and spatial structure in a wild bird population.” American Naturalist. 10.1086/738329

74. Zanette, Liana Y., Aija F. White, Marek C. Allen, and Michael Clinchy. 2011. “Perceived Predation Risk Reduces the Number of Offspring Songbirds Produce per Year.” Science 334 (6061): 1398–1401. 10.1126/science.1210908

